# ALS Motor Neurons Exhibit Hallmark Metabolic Defects That Are Rescued by Nicotinamide and SIRT3 Activation

**DOI:** 10.1101/713651

**Authors:** Jin-Hui Hor, Munirah Mohamad Santosa, Valerie Jing Wen Lim, Beatrice Xuan Ho, Amy Taylor, Zi Jian Khong, John Ravits, Yong Fan, Yih-Cherng Liou, Boon-Seng Soh, Shi-Yan Ng

**Author notes:** Correspondence to: Shi-Yan Ng, Boon-Seng Soh.

## Abstract

Motor neurons (MNs) are highly energetic cells and recent studies suggest that altered energy metabolism precede MN loss in Amyotrophic Lateral Sclerosis (ALS), an age-onset neurodegenerative disease. However, clear mechanistic insights linking altered metabolism and MN death are still missing. In this study, induced pluripotent stem cells (iPSCs) from healthy controls, familial ALS and sporadic ALS patients were differentiated towards spinal MNs, cortical neurons and cardiomyocytes. Metabolic flux analyses reveal a MN-specific deficiency in mitochondrial respiration in ALS. Intriguingly, all forms of familial and sporadic ALS MNs tested in our study exhibited similar defective metabolic profiles, which were attributed to hyper-acetylation of mitochondrial proteins. In the mitochondria, SIRT3 functions as a mitochondrial deacetylase to maintain mitochondrial function and integrity. We found that activating SIRT3 using nicotinamide or a small molecule activator reversed the defective metabolic profiles in all our ALS MNs, as well as correct a constellation of ALS-associated phenotypes.

## INTRODUCTION

Amyotrophic Lateral Sclerosis (ALS) is an age-onset, progressive neurodegenerative disorder affecting both upper and lower motor neurons (MNs). The loss of MNs leads to denervation of skeletal muscles, thereafter, leading to stiff muscles and muscular atrophy that results in difficulties in speech, swallowing, walking and breathing. Majority of ALS cases (up to 90%) are sporadic, where the cause of disease is largely unknown. The rest of ALS patients have a familial form of the disease where mutations in genes such as SOD1, C9ORF72 and TDP43 are most common. Despite the genetic differences, the clinical manifestations of sporadic and familial ALS patients are indiscernible, suggesting a possible converging pathogenic mechanism.

In patients, besides motor neuron degeneration, ALS is associated with impairments of energy metabolism, and clinical evidence supports a positive correlation between defective energy metabolism and rate of ALS progression ^1^. Animal models of ALS also support metabolic dysregulation as a contributing pathogenic pathway. Previous studies reported that SOD1^G93A^ mice developed progressive central nervous system acidosis as ALS advanced, with the lowest pH recorded in spinal cords of end-stage and moribund mice ^2^, suggesting that neural cell metabolism in ALS is affected.

Increasingly, studies have shown that mitochondrial morphology abnormalities are associated with both familial and sporadic ALS ^3–5^. It has been suggested that these abnormalities impair oxidative phosphorylation, contributing to the ALS pathology ^6^. Mitophagy, a process where damaged mitochondria is selectively degraded by autophagy, has also been shown to be elevated in SOD1 mutant mice, and that halting mitophagy in these mice delayed ALS progression and improved survival ^7^. Collectively, these evidences suggest that mitochondrial dysfunctions contribute to ALS pathogenesis.

Neurons rely mainly on oxidative phosphorylation to fuel their high metabolic demands, and any deviation from this norm would lead to neurological disorders ^8,9^. In this current work, we investigated if defective mitochondrial respiration could be the common pathway implicated in both sporadic and familial ALS. To do so, we differentiated patient-derived and isogenic pairs of induced pluripotent stem cells (iPSCs) towards spinal MNs and developed a method to enrich for these MNs for metabolic flux measurements. By excluding neural progenitor cells (NPCs) refractory to differentiation and other non-neuronal cells in our metabolic assays, we identified reduced mitochondrial respiration and elevated glycolysis as a metabolic hallmark of ALS MNs, which were not observed in the NPCs. We found that the defective mitochondrial respiration was attributed to the hyper-acetylation of mitochondrial proteins caused by reduction of SIRT3 activity. Using nicotinamide and small molecule activators to restore SIRT3 activity, we were not only able to rescue the metabolic defects, but also improve ALS MN morphology and promoted survival. Our work demonstrates that hyper-acetylated mitochondrial proteins is a hallmark of both sporadic and familial ALS and elevating SIRT3 activity can be explored as a therapeutic strategy for ALS.

## RESULTS

### MNs generated from ALS patient iPSCs and isogenic ALS knock-in iPSCs exhibit reduced mitochondrial respiration and ATP production

MNs were differentiated from 3 healthy iPSC lines: BJ-iPS, 18a and GM23720, 3 sporadic ALS lines: sALS1, sALS2 and sALS3, as well as familial ALS iPSCs (harboring the following mutations): 29d (SOD1^L144F^), 49a (TDP43^G298S^) and 19f (C9ORF72 expanded GGGGCC repeats) ^10^ using established protocols (**Fig. 1a**). Using this chemically defined protocol, ISL1^+^SMI32^+^ MNs were effectively derived from all of these iPSC lines by day 28 (**Fig. 1b**). To confirm ALS phenotypes in these cell lines that were previously reported, we measured basal survival of ISL1^+^ MNs from day 25 to day 35 and found that an accelerated death phenotype is associated with the ALS MNs (**Fig. 1c**). Since elevated ER stress is also a molecular signature of ALS MNs ^11^, we also measured mRNA levels of key genes in the ER stress pathway and found significant upregulation of *CHOP* and spliced XBP1 (*sXBP1*) (**Fig. 1d**).

**Figure 1:**
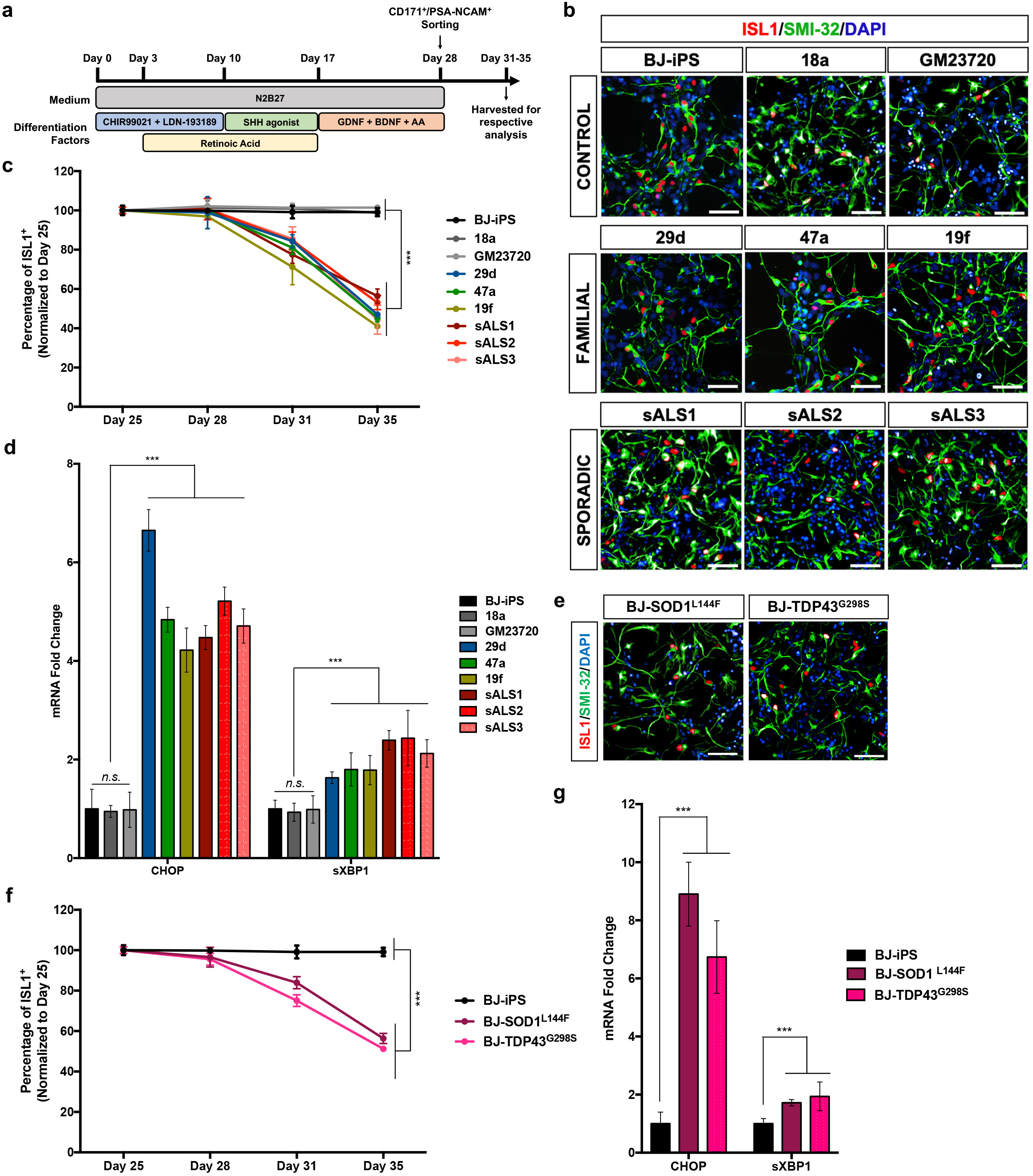
ALS iPSC-derived MNs exhibit diseased phenotypes. **(a)** Schematic of the MN differentiation protocol. **(b)** Immunostaining of wild-type (BJ-iPS, 18a and GM23720), familial ALS (29d, 47a and 19f) and sporadic ALS (sALS1, sALS2, sALS3) iPSC-derived cultures at day 28 indicating the derivation of ISL1^+^SMI32^+^ MNs. Cellular nuclei were counterstained with DAPI. Scale bars, 50 μm. **(c)** Quantification of ISL1^+^ MNs from day 25 to day 35 demonstrating that wild-type MNs (BJ-iPS, 18a and GM23720) remain viable while familial and sporadic ALS MNs show significantly reduced survival over time. **(d)** qPCR quantification of ER stress transcripts *CHOP* and spliced *XBP1* (*sXBP1*) in MN cultures at day 28. Fold changes are normalized to expression levels of respective mRNA in BJ-iPS. **(e)** Immunostaining of isogenic ALS (BJ-SOD1^L144F^ and BJ-TDP43^G298S^) iPSC-derived cultures at day 28 indicating the formation of ISL1^+^SMI32^+^ MNs. Cellular nuclei were counterstained with DAPI. Scale bars, 50 μm. **(f)** Quantification of ISL1^+^ MNs derived from BJ-SOD1^L144F^ and BJ-TDP43^G298S^ from day 25 to day 35 revealed an accelerated death phenotype similar to that of other ALS lines. **(g)** MN cultures derived from BJ-SOD1^L144F^ and BJ-TDP43^G298S^ show upregulation of *CHOP* and *sXBP1* compared to its isogenic control line BJ-iPS. In **(d)** and **(g)**, gene expression was normalized to ACTINB and HPRT. ***p < 0.001, n.s. non-significant; two-tailed t test.

To ensure that the ALS-specific phenotypes and molecular profiles are not due to inherent variability between cell lines, we generated isogenic controls where we introduced the SOD1^L144F^ and TDP43^G298S^ mutations into the healthy BJ-iPS line using the CRISPR/Cas9 technology (**Fig. 1e**; **Extended Data Figs. 1a, 1b**). Mutations were then confirmed by DNA-sequencing (**Extended Data Figs. 1a, 1b**). Similar to what was observed with the patient iPSC lines, MNs derived from the isogenic SOD1^L144F^ (BJ-SOD1^L144F^) and TDP43^G298S^ (BJ-TDP43^G298S^) knock-in lines also showed decreased MN survival and elevated expression of ER stress genes *CHOP* and *sXBP1* (**Figs. 1f, 1g**).

Although previous studies showed that ALS neurons have abnormal mitochondria ^3,4^, it has not been established if these morphological abnormalities have any effects on mitochondrial functions in ALS MNs. Since oxidative phosphorylation is critical for maintenance of neuronal metabolism and survival, we investigated if mitochondrial respiration in ALS MNs could be compromised. To enrich for MNs in our iPSC-derived cultures, we performed magnetic sorting using a cocktail of PSA-NCAM and CD171 antibodies. Using this sorting strategy, we enriched for ISL1^+^ MNs to approximately 60% (**Figs. 2a, 2b**) without the use of AraC, which is sometimes used to deplete neural progenitor cells (NPCs) in the cultures but also induces neuronal death through oxidative stress ^12^. Oxygen consumption rate (OCR) of these sorted neurons was measured as a function of time using an extracellular flux analyzer. The MitoStress Assay was used to measure bioenergetics parameters, which revealed lower basal respiration, ATP production and maximal capacity in all familial and sporadic ALS lines compared to healthy MNs (**Figs. 2c, 2d**). Likewise, MNs derived from BJ-SOD1^L144F^ and BJ-TDP43^G298S^ isogenic iPSCs exhibited similar reductions in basal respiration (*p* < 0.0001), ATP production (*p* < 0.0001) and maximal capacity (*p* < 0.01), similar to 29d and 47a MNs that also possess the heterozygous L144F mutation in SOD1 and G298S mutation in TDP43 respectively (**Figs. 2e, 2f**).

**Figure 2:**
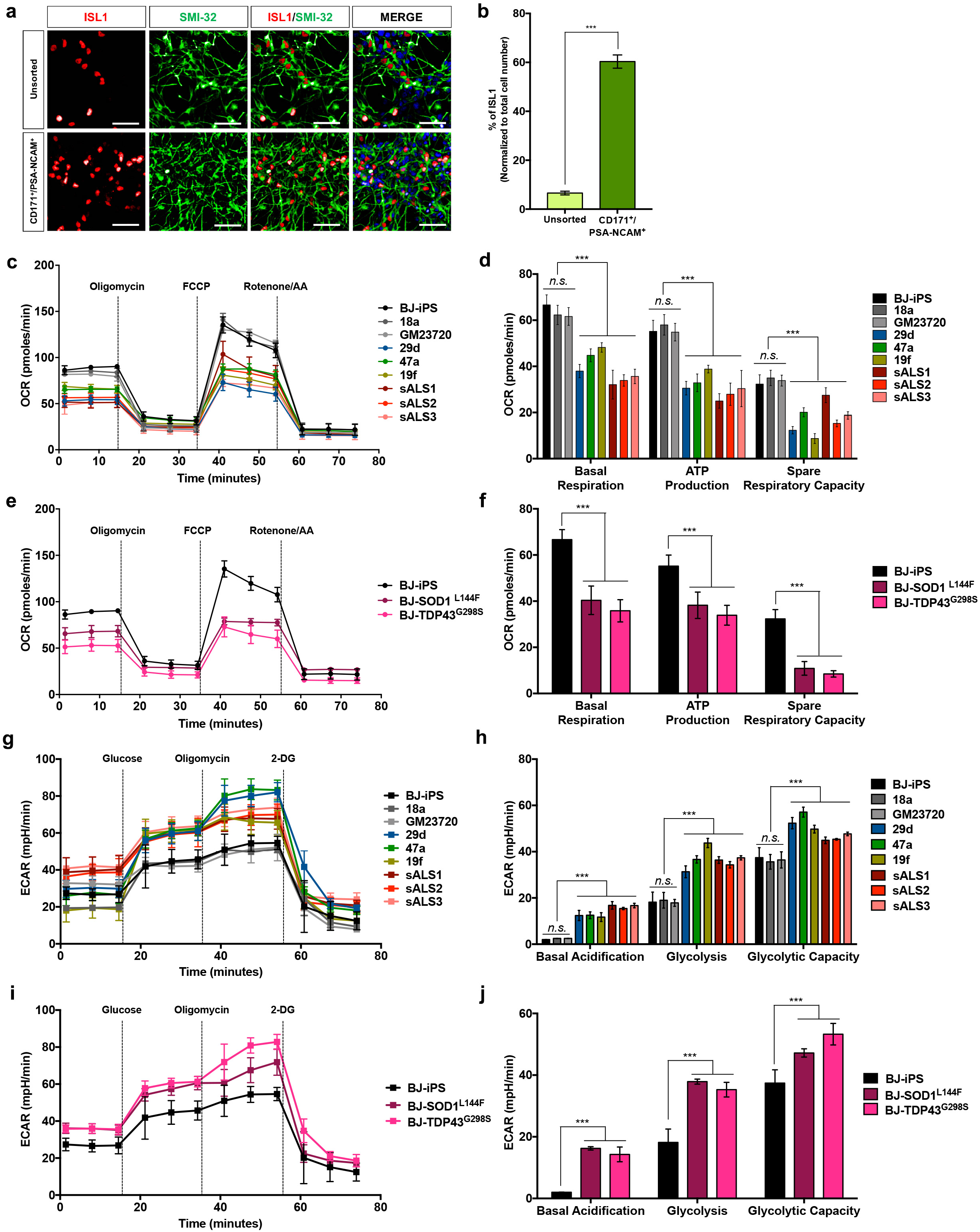
Sporadic and familial ALS MNs show a hypo-oxidative and hyper-glycolytic metabolic profile. **(a)** Immunostaining of unsorted cultures at day 28 and cultures sorted with CD171 and PSA-NCAM with ISL1 and SMI32 demonstrating enrichment of ISL1^+^ MNs in the sorted cultures. Cellular nuclei were counterstained with DAPI. Scale bars, 50 μm. **(b)** Quantification of ISL1^+^ MN numbers in unsorted and sorted cultures at day 28 indicating more than 5-fold enrichment (from 7% to 59%). **(c)** Metabolic flux plots of healthy and ALS patient-derived sorted neurons, where oxygen consumption rate (OCR) was measured as a function of time. The MitoStress assay was used to measure bioenergetics parameters, by adding Complex V inhibitor Oligomycin, mitochondrial uncoupler FCCP and Complexes I and III inhibitors Rotenone and Antimycin A (AA). **(d)** Basal respiration, ATP production and spare respiration were calculated for sorted neurons from each of the cell lines. **(e-f)** OCR measurements using the MitoStress assay were performed and calculated for the isogenic pairs BJ-iPS, BJ-SOD1^L144F^ and BJ-TDP43^G298S^. **(g)** Metabolic flux plots of healthy and ALS patient-derived sorted neurons, where extracellular acidification rate (ECAR) was measured as a function of time. The Glycolysis stress assay was used to measure bioenergetics parameters, by adding glucose, Complex V inhibitor Oligomycin, and hexokinase inhibitor 2-DG. **(h)** Basal acidification, glycolysis and glycolytic capacity were calculated for sorted neurons from each of the cell lines. **(i-j)** ECAR measurements using the Glycolysis stress assay was performed and calculated for the isogenic pairs BJ-iPS, BJ-SOD1^L144F^ and BJ-TDP43^G298S^. ***p < 0.001, n.s. non-significant; two-tailed t test.

Since ATP production in diseased MNs is reduced by at least 25% in the various ALS lines, we reasoned that these neurons would have to turn to other sources of energy to fuel their metabolic needs. Glycolysis, measured by extracellular acidification rate (ECAR), revealed that the ALS MNs exhibited increased glycolysis and glycolytic capacity (**Figs. 2g, 2h**). Likewise, the elevated glycolytic profiles in BJ-SOD1^L144F^ and BJ-TDP43^G298S^ MNs compared to BJ-iPS MNs confirmed that this is an ALS-specific metabolic feature rather than due to variation between cell lines (**Figs. 2i, 2j**). Importantly, these metabolic changes are specific to the MN stage. OCR measurements of day 10 NPCs revealed no significant changes to the basal respiration and ATP production in the ALS cells, even though the total capacity of ALS NPCs was approximately halved (**Extended Data Figs. 2a-2d**). Furthermore, glycolysis profiles of day 10 NPCs do not show significant changes between healthy and ALS cells (**Extended Data Figs. 2e-2h**).

### Other metabolically active cell types derived from ALS iPSCs do not show overt reduction in mitochondrial respiration

Next, we wondered if ALS-causing mutations affect all actively respiring cell types or specifically MNs. To this end, we differentiated all iPSCs towards cTnT^+^ cardiomyocytes (**Extended Data Fig. 3a**) and metabolic flux analyses were performed. BJ-SOD1^L144F^ and BJ-TDP43^G298S^ cardiomyocytes were not significantly different from their isogenic healthy control in terms of basal respiration and ATP production, although total respiratory capacity was reduced (**Extended Data Fig. 3b**). Likewise, similar results were also obtained for familial and sporadic ALS patient-derived cardiomyocytes (**Extended Data Fig. 3c**). Taken together, our results revealed that reduced mitochondrial respiration and the concomitant elevation of glycolysis is a metabolic hallmark of ALS MNs.

### ALS MN metabolic defects caused by hyper-acetylation of mitochondrial proteins

Having established that mitochondrial respiration defects are hallmark of both familial and sporadic ALS, we sought to understand the mechanisms that underlie these defects, and whether correction of this metabolic defect would alleviate ALS phenotypes. Mitochondrial respiration is known to be fine-tuned by the acetylation of mitochondrial proteins, and this process is controlled by the mitochondrial deacetylase SIRT3 ^13^. Previous studies have established that SIRT3 depletion leads to mitochondrial protein hyper-acetylation in muscles, cardiac and liver tissues, along with dysregulation of mitochondrial respiration (refs). Therefore, we investigated if the loss of SIRT3 in MNs would result in the ALS-specific metabolic defects we observed. Western blot analysis, however, did not reveal significant changes in SIRT3 levels between healthy and ALS MNs or MNs from the set of isogenic iPSC lines (**Figs. 3a, 3b**). We postulate that SIRT3 activity could be affected even though expression levels were unchanged. Therefore, to determine SIRT3 activity, we measured relative acetylation of lysine 68 on MnSOD (MnSOD K68ac), one of the best-characterized SIRT3 target ^14,15^. Indeed, Western blot analyses revealed significantly higher MnSOD K68ac signals in all of our ALS MN cultures, including the sporadic ALS MNs and isogenic MNs, compared to healthy controls, suggesting reduced SIRT3 activity (**Figs. 3a, 3c**). Human post-mortem lumbar spinal cord sections were also analyzed by immunohistochemistry, which confirmed higher MnSOD K68ac signals in alpha MNs of lumbar spinal cord in sporadic ALS patients compared with non-ALS controls (**Figs. 3d, 3e**). Since SIRT3 has multiple targets in the mitochondria, it is expected that loss of SIRT3 activity would impact global mitochondrial acetylation. To confirm this, mitochondrial extracts from all the iPSC-derived MNs were analyzed and immunoblotting was performed using a specific antibody against acetyl-lysine residues. Western blot analysis revealed significantly higher intensities of acetylated proteins in all of the ALS lines compared to the healthy controls, concurring with our MnSOD K68ac assay (**Figs. 3a, 3f**). Previous SIRT3 interactome studies have revealed several Complexes I subunits amongst its downstream targets ^16^. Expanding on these studies, we found that reduced Complex I activity was observed in all of our ALS MNs (**Fig. 3g**), which explained the lower basal mitochondrial respiration observed in these ALS MNs (**Fig. 2**).

**Figure 3:**
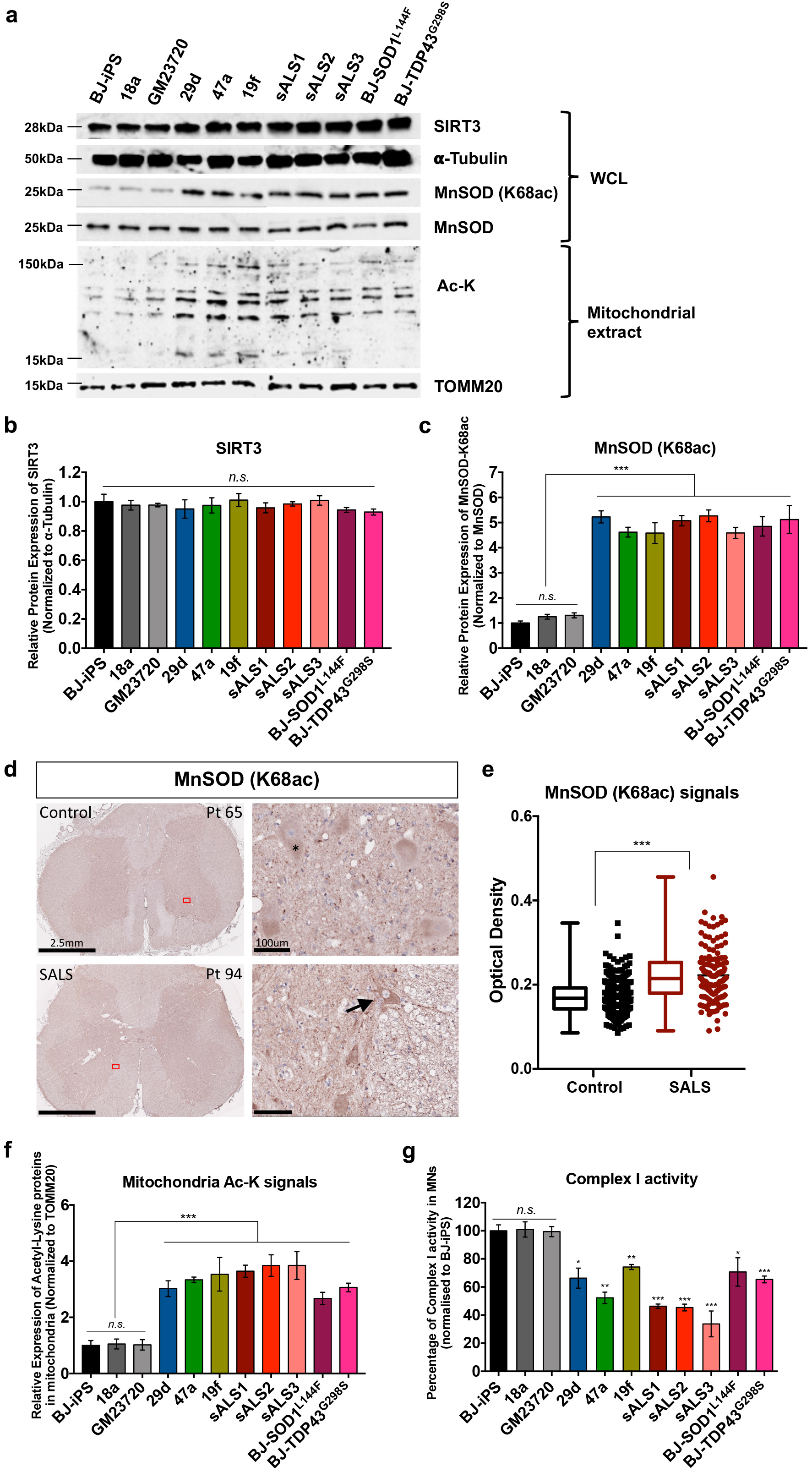
Hyper-acetylation of mitochondrial proteins in familial and sporadic ALS MNs. **(a)** Western blot analyses of iPSC-derived MNs at day 28 probing for SIRT3, total MnSOD, MnSOD with specific acetylation at lysine-68 (MnSOD K68ac) on whole cell lysate as well as probing for acetyl-lysine proteins in purified mitochondrial extracts. **(b)** Densitometric analyses of Western blot bands reveal no significant changes in SIRT3 protein levels in ALS versus healthy MNs. **(c)** Densitometric analysis of MnSOD(K68ac) normalized to total MnSOD revealed approximately -fold upregulation of MnSOD(K68ac) in all of our ALS iPSC-derived MNs. This indicates reduced SIRT3 activity in ALS MNs. **(d)** Immunohistochemistry of control and sporadic ALS patients (SALS) lumbar sections revealed increased MnSOD (K68ac) signals in SALS patients lumbar motor neurons (arrow). Lipofuscin is visible in large, healthy motor neurons, a function of normal cellular aging and unrelated to disease (*). Scale bars, 2.5mm (left panel) and 100 μm (right panel). **(e)** Quantification of MnSOD (K68ac) signals in both control (n=380) and SALS (n=216) lumbar motor neurons demonstrates increased MnSOD (K68ac) signals in SALS patients. **(f)** Densitometric analysis of acetyl-lysine signals normalized to TOMM20 revealed approximately 3-fold increase in acetylation of mitochondria proteins in all of our ALS iPSC-derived MNs. **(g)** Complex I activity in healthy and ALS MNs were measured, which revealed significant decline of between 30 to 80% in the ALS MNs. *p<0.05, **p<0.01, ***p < 0.001, n.s. non-significant; two-tailed t test.

### Loss of SIRT3 function mimics ALS phenotypes

To confirm that reduced SIRT3 activity is responsible for the mitochondrial respiration defects, we created genetic models of SIRT3 depletion and performed RNA interference to study effects of SIRT3 depletion on MN survival and function. Using the CRISPR/Cas9 approach, we generated multiple isogenic SIRT3 haploinsufficient (SIRT3^+/−^) iPSC lines (**Extended Data Fig. 1c**). Out of 66 clones that were screened in total, 31 were heterozygous knockouts (47%) while none of them were total knockouts, suggesting that SIRT3-deficient iPSCs were unviable. We randomly selected SIRT3^+/−^ clones #6 and #17 and showed that both clones differentiated into MNs with approximately the same efficiencies as the isogenic control BJ-iPS (**Extended Data Fig. 1d**). We verified both BJ-SIRT3^+/−^ #6 and #17 by Western blot that showed a 50% reduction of SIRT3 protein. MnSOD K68ac levels were also significantly higher in BJ-SIRT3^+/−^ #6 and #17, confirming reduced SIRT3 activity (**Figs. 4a, 4b**). Additionally, purified mitochondrial extracts also showed that clones #6 and #17 had more acetyl-lysine residues (**Fig. 4a**), consistent with the hyper-acetylated mitochondrial profiles seen in the ALS iPSC-derived MNs (**Fig. 3a**). A constellation of *in vitro* ALS phenotypes has now been reported, which includes accelerated MN death, elevated ER stress signaling, smaller cell bodies ^11,17^, and a hypo-oxidative/hyper-glycolytic metabolic profile (**Fig. 2**). Therefore, we sought to determine if loss of SIRT3 could induce ALS phenotypes *in vitro*. Measurements of ER stress transcripts in the day 28 MNs from both SIRT3^+/−^ clones revealed significant upregulation of *CHOP* and *sXBP1* mRNAs (**Fig. 4c**), similar to that seen in all the ALS MNs we tested (**Fig. 1d**). Metabolic flux measurements confirmed that MNs derived from both SIRT3^+/−^ clones exhibited reduced mitochondrial respiration (**Figs. 4d, 4e**) and elevated glycolysis simultaneously (**Figs. 4f, 4g**). Phenotypically, motor neurons derived from both SIRT3^+/−^ clones demonstrated reduced survival (**Fig. 4h**) and significantly reduced soma sizes and primary neurites at day 31 (**Figs. 4i, 4j**). Given that SIRT3^+/−^ MNs already display ALS-like phenotypes, we are convinced that even partial loss of SIRT3 activity contributes to ALS pathogenesis.

**Figure 4:**
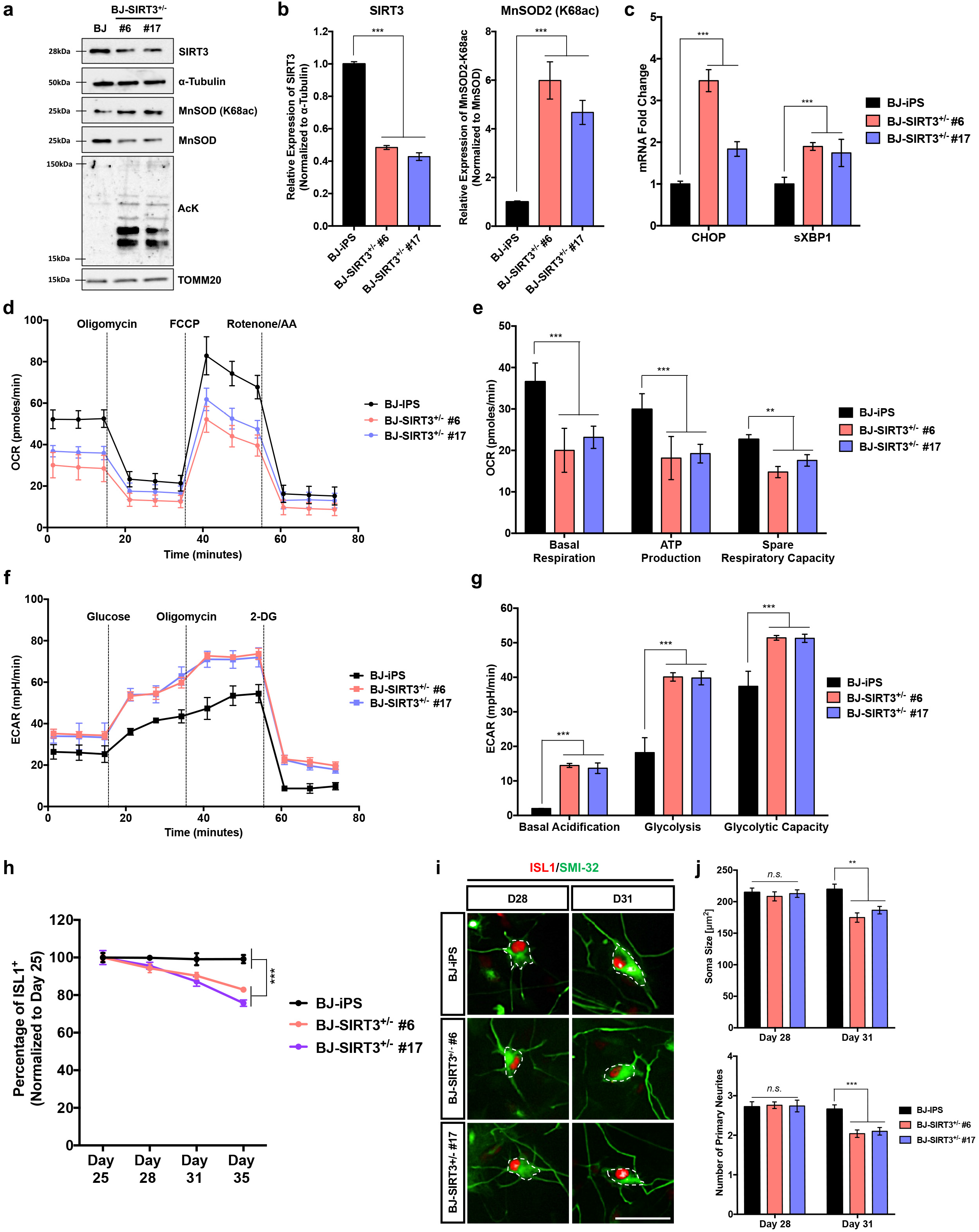
SIRT3 deficiency in MNs results in ALS-like phenotypes. **(a)** Western blot analysis of day 28 MNs derived from BJ-iPS and two isogenic SIRT3^+/−^ (#6 and #17) clones confirmed reduction in SIRT3 protein, along with increased MnSOD (K68ac) and increased acetylation of mitochondrial proteins. **(b)** Densitometric analyses of Western blot bands reveal 50% decrease in SIRT3 protein levels and increased MnSOD (K68ac) in both SIRT3^+/−^ #6 and #17 versus healthy MNs. **(c)** qPCR measurements of *CHOP* and *sXBP1* show significant upregulation of both ER stress transcripts in SIRT3^+/−^ #6 and #17 relative to the isogenic BJ-iPS control. **(d)** Measurements of OCR using the MitoStress assay of day 28 MNs differentiated from BJ-iPS (shown in black), SIRT3^+/−^ #6 (pink) and #17 (violet). **(e)** Measurements of basal respiration, ATP production and spare respiration of day 28 MNs differentiated from BJ-iPS (shown in black), SIRT3^+/−^ #6 (pink) and #17 (violet). **(f)** Measurements of ECAR using the Glycolysis Stress assay of day 28 MNs differentiated from BJ-iPS (shown in black), SIRT3^+/−^ #6 (pink) and #17 (violet). **(g)** Measurements of basal acidification, glycolysis and glycolytic capacity of day 28 MNs differentiated from BJ-iPS (shown in black), SIRT3^+/−^ #6 (pink) and #17 (violet). **(h)** Quantification of ISL1^+^ MNs derived from both BJ-SIRT3^+/−^ clones from day 25 to day 35 revealed a progressive death phenotype as compared to its healthy control (BJ-iPS). **(i)** Representative images of ISL1^+^SMI32^+^ MNs derived from BJ-iPS, BJ-SIRT3^+/−^ #6 and #17 iPSCs, showing cell body sizes (outlined in white dotted lines) from MNs at day 28, and at day 31. Scale bars, 50 μm. **(j)** Quantification of mean cell body size and number of primary neurites of both BJ-SIRT3^+/−^ clones reveal deteriorating neuronal health from day 28 to day 31. **p<0.01, ***p < 0.001, n.s. non-significant; two-tailed t test.

To confirm our earlier findings with SIRT3^+/−^ MNs, we performed small interfering RNA (siRNA)-mediated knockdown to transiently deplete SIRT3 levels in healthy BJ-iPS MNs at day 25. At day 28, we confirmed 70% knockdown at both mRNA and protein levels, together with increased MnSOD K68ac, indicative of decreased SIRT3 activity (**Extended Data Figs. 4a, 4b**). Analyses of these neurons at day 28 also revealed reduced OCR parameters (**Extended Data Fig. 4c**). This set of siRNA experiments confirms that SIRT3 depletion in MNs is sufficient to cause the ALS-like phenotypes.

### Nicotinamide supplementation and SIRT3 activation reverse ALS MN phenotypes

Since SIRT3 is a NAD^+^-dependent deacetylase, one possibility for reduction in SIRT3 activity could be insufficient levels of intracellular NAD^+^. To investigate this, we measured NAD^+^ levels in our ALS MNs, and found 20-30% reduction in NAD^+^ levels in all of the ALS MNs (**Fig. 5a**). Therefore, we investigated the effects of NAD^+^ supplementation on ALS MNs. Following previous published work, we supplemented MN cultures with 0.5 mM Nicotinamide (NAM), a precursor of NAD^+^, from days 28 to 31, and performed the ALS phenotypic and metabolic assays. Compared to the water control, NAM supplementation promoted sporadic and familial ALS MN survival (**Fig. 5b**) and morphologies (**Extended Data Fig. 6a, 6b**). Additionally, basal mitochondrial respiration, ATP production as well as spare respiratory capacity were significantly improved (**Figs. 5c-5e**). It is worth pointing out that NAM supplementation had no significant effects on healthy MNs, indicating targeted rescue of ALS MNs.

**Figure 5:**
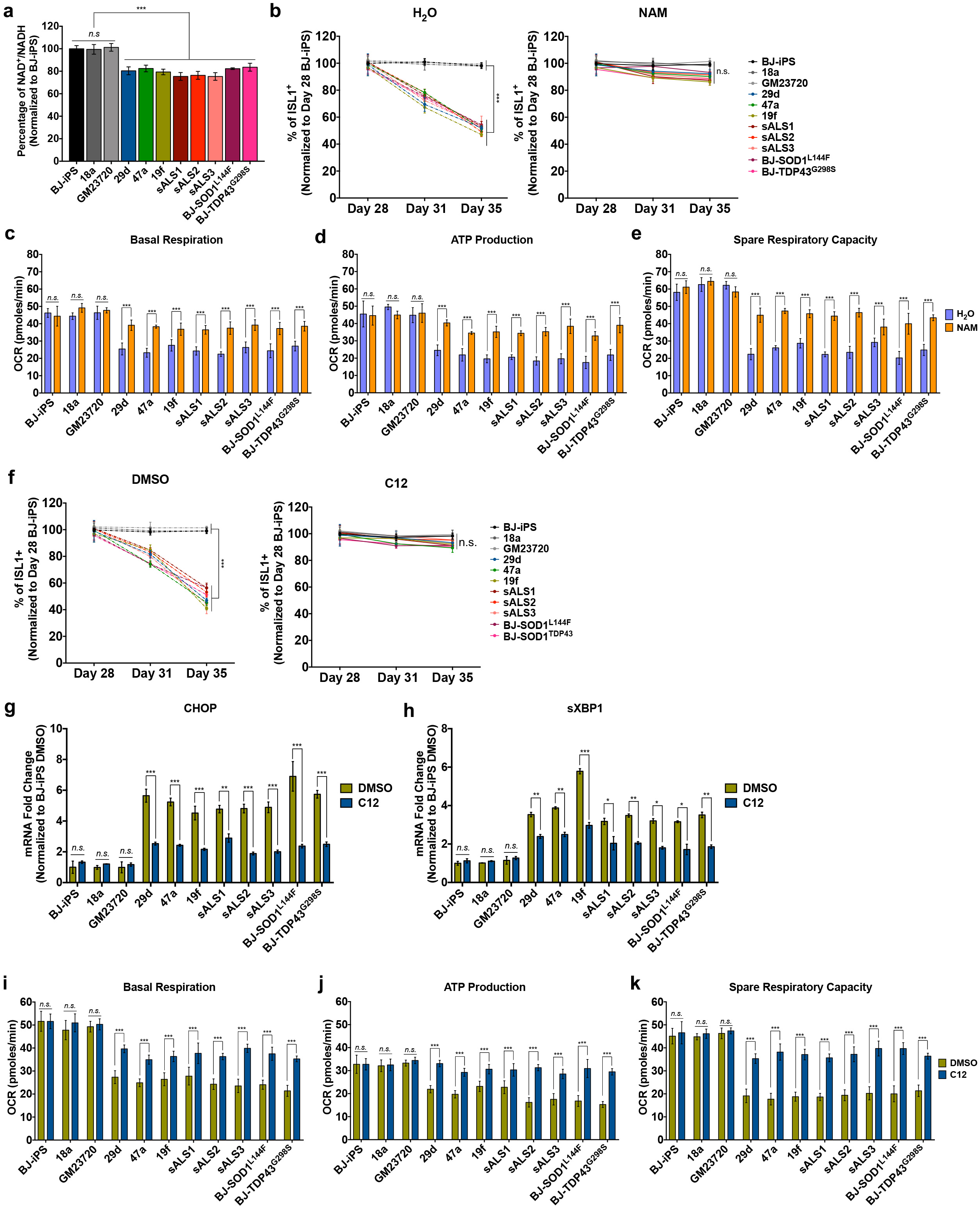
Nicotinamide treatment or SIRT3 activation reverses mitochondrial respiration defect in ALS MNs. **(a)** Measurements of intracellular NAD+:NADH ratio in healthy and ALS MNs revealed reduced NAD+ availability in ALS MNs. **(b)** WT and ALS iPSC-derived MNs were treated with either H_2_O or NAM from day 28 to day 35. Number of ISL1^+^ MNs were quantified and normalized to number of ISL1^+^ MNs in respective cell lines at day 28. NAM supplementation prevents MN death in ALS MNs. **(c-e)** Measurements of basal respiration, ATP production and spare respiration respectively in healthy and ALS MNs treated with water as a control (purple) or 0.5 mM NAM (orange). **(f)** WT and ALS iPSC-derived MNs were treated with either DMSO or C12 from day 28 to day 35. Number of ISL1^+^ MNs were quantified and normalized to number of ISL1^+^ MNs in respective cell lines at day 28. C12 treatment prevents MN death in ALS MNs. **(g-h)** qPCR quantification of ER stress transcripts CHOP and spliced XBP1 (sXBP1) in MN cultures at day 31 treated with DMSO or C12. Fold changes are normalized to expression levels of respective mRNA in BJ-iPS MNs treated with DMSO. **(i-k)** Measurements of basal respiration, ATP production and spare respiration respectively in healthy and ALS MNs treated with DMSO (green) and C12 (blue). Of note, C12 treatment completely rescued ATP production back to healthy levels. ***p < 0.001, n.s. non-significant; two-tailed t test.

Next, we wondered if overexpression of SIRT3 could rescue ALS MN phenotypes. Using either an inducible SIRT3-expressing lentivirus or control GFP virus, we infected all our iPSC-derived MNs at day 28 and harvested them at day 31 for analyses. Immunoblotting confirmed that SIRT3 is reliably over-expressed in all our 11 iPSC-derived MNs, but MnSOD K68ac signals were not reduced in the SIRT3 over-expressing MNs (**Extended Data Fig. 4d**). We also noted that SIRT3 overexpression significantly improved mitochondrial respiration in the three healthy lines but has no significant effects in all the ALS MNs (**Extended Data Figs. 4e-4g**), confirming our earlier results that SIRT3 activity rather than expression is limiting in the ALS MNs. Therefore, it is not surprising that simply overexpressing SIRT3 without the addition of co-factors would have no significant effects on mitochondrial respiration in ALS MNs.

A small number of SIRT3 activators have been identified to date, including a specific SIRT3 agonist previously identified as 7-hydroxy-3-(4’-methoxyphenyl) coumarin or C12 that binds to SIRT3 with high affinity and promotes deacetylation of downstream targets ^15^. Through a dose-response test, we determined that 5 μM C12 was able to reduce MnSOD K68ac signals without cytotoxicity (**Extended Data Figs. 5a-5d**). Treatment of all our iPSC-derived MNs with 5 μM C12 from days 28 to 31 confirmed that mitochondrial acetyl-lysine residues signals were significantly reduced in the ALS MNs (**Extended Data Fig. 5e**). Consequently, Complex I activity of ALS MNs significantly improved after 3 days of C12 treatment (**Extended Data Fig. 5f**). To investigate effects of C12 treatment on ALS MNs, we treated our repertoire of iPSC-derived MNs with either C12 or DMSO control from day 28 to 31. Similar to our observations for NAM, we found that C12 promoted ALS MN survival (**Fig. 5f**) and significantly reduced expressions of ER stress transcripts *CHOP* and *sXBP1* in all the ALS MNs (**Fig. 5g, 5h**). Metabolic flux analyses revealed that C12 was able to promote mitochondrial respiration (**Figs. 5i-k**) and lower glycolysis (**Extended Data Fig. 5g-i)** in ALS MNs. Finally, neuronal morphologies were improved with C12 treatment, resulting in enlarged soma sizes and increased number of primary neurites similar to healthy MNs (**Extended Data Fig. 6c, 6d**), similar to our observations with NAM.

Notably, treatment of BJ-SIRT3^+/−^ #6 and #17 with C12 did not promote survival or improved neuronal morphology (**Extended Data Figs. 7a, 7b**). C12 treatment on both SIRT3-deficient clones also did not improve mitochondrial respiration (**Extended Data Fig. 7c**), providing additional evidence that C12 works through promoting SIRT3 activity rather than via non-specific off-target effects.

## DISCUSSION

ALS is a heterogeneous motor neuron disease, yet all patients have similar clinical manifestations, suggesting the possibility of an upstream pathogenic pathway independent of the various genetic mutations known to cause ALS. In this study, we identified a metabolic hallmark of both sporadic and familial ALS MNs that is characterized by hypo-oxidation and hyper-glycolysis. Increased production and release of lactate, the end product of the glycolytic pathway, would explain the central nervous system acidosis reported in SOD1^G93A^ mice ^2^. It is highly possible that the hyper-glycolytic metabolism was a compensatory mechanism to overcome the lack of ATP generated through oxidative phosphorylation in ALS MNs, because restoration of ATP production through NAM supplementation and SIRT3 activation also corrected the hyper-glycolytic metabolic profile (**Fig. 5**). In addition to using patient-derived iPSCs, isogenic cell lines were made to confirm our findings that loss of SIRT3 activity was responsible for the mitochondrial metabolic defects, and the constellation of ALS-like phenotypes. We note that we were unable to obtain SIRT3^−/−^ iPSCs in this study, while mice with complete Sirt3-knockout develop normally and are not embryonic lethal, although neuronal survival and function appear to be affected ^18^. We believe these could be due to species-specific differences and are carrying out additional investigation in a separate study. Nevertheless, our results confirmed that partial loss of SIRT3 was sufficient to cause ALS phenotypes in iPSC-derived MNs.

One of our key findings was that reduced SIRT3 activity contributed to mitochondrial hyper-acetylation of mitochondrial proteins and subsequent metabolic defects that are novel hallmarks of ALS MNs. MNs derived from familial and sporadic patient iPSCs, as well as isogenic iPSC lines with SOD1^L144F^ and TDP43^G298S^ mutations demonstrate a consistent increase in acetylated mitochondrial proteins, including MnSOD K68ac, a well characterized target of SIRT3. Post-mortem analyses of ALS spinal cords further corroborate this finding. Importantly, elevating SIRT3 activity using either Nicotinamide or C12 reversed multiple ALS MN defects.

Our data supports recently published work that NAD^+^ levels in ALS patients are reduced compared to the healthy population ^19^, and confirms that supplementation of NAM, a precursor of NAD^+^, reverses ALS phenotypes. Lack of NAD^+^ has also been previously implicated in MN dysfunction where intracellular nicotinamide phosphoribosyltransferase (iNAMPT) knockout in adult mice results in a MN degeneration phenotype mimicking ALS ^20^. iNAMPT functions as the rate-limiting enzyme of the mammalian NAD^+^ biosynthesis salvage pathway. Taken together, these independent studies also suggest that MNs are particularly sensitive to NAD^+^ reductions.

In conclusion, our study using patient-derived and isogenic iPSCs reveals that reduced mitochondrial respiration and elevated glycolysis are metabolic hallmarks of ALS MNs. We have also established that activation of mitochondrial SIRT3 is a target for reversing disease phenotypes for both sporadic and familial ALS. Finally, our data confirms that NAM supplementation, as well as a small molecule SIRT3 agonist reverse several *in vitro* ALS phenotypes and has therapeutic potential for development into an effective treatment.

## METHODS

### Human iPSC culture and differentiation towards MNs and cardiomyoctes

Human iPSCs were cultured feeder-free on Matrigel-coated dishes with iPS-Brew (Miltenyi Biotec). The iPSC lines, and their mutations, used in this study are listed in Supplementary Table S1. To passage the iPSCs, confluent iPSCs were sub-cultured using ReLeSR (Stem Cell Technologies) following the manufacturer’s instructions. Pluripotent stem cells were differentiated towards the spinal motor neuron fate following established protocols described previously^21^. Pluripotent stem cells were differentiated towards the cardiomyocytes fate following established protocols described previously^22^.

### CRISPR/Cas9-mediated genome editing

Guide RNAs (gRNAs) were designed using Feng Zhang lab’s guide design tool at crispr.mit.edu before it shut down: gRNAs were cloned into a Cas9-containing plasmid PX458 or PX459 and transfected into 293T cells using Lipofectamine 2000 Transfection Reagent for surveyor nuclease assay. Verified gRNAs and ssODNs (Supplementary Table S2) were transfected into BJ-iPS hiPSCs using Lipofectamine Stem Transfection Reagent. 2 days after transfection, cultures were sorted for GFP+ cells or selected with 1 μM puromycin. Single cells were then plated out and allowed to expand before screening. gDNA of the colonies were collected and subjected to PCR amplification before sanger sequencing (Applied Biosystems 3730xl).

### Survival curve for motor neuron cultures

Motor neurons was treated with AraC on Day 23 and plated at 75,000 cells per well of a 96 well plate on Day 24. Cultures were then 4% PFA on Day 25, 28, 31 and 35 for quantification of motor neurons survival. Biological triplicates were performed with a minimum of 5 technical replicates each.

### Magnetic microbead sorting of neurons

After dissociating with Accutase, cells were blocked with solution containing phosphate-buffered saline, 0.5% bovine serum albumin (BSA) and 2mM EDTA. The cells were then incubated with CD171-APC antibody (Miltenyi Biotec) and PSA-NCAM-APC antibody (Miltenyi Biotec) for 10 mins at 4°C. After washing, the cells were then incubated with anti-APC microbeads for 15 minutes at 4°C. The cells were then washed twice and filtered prior to loading into the separation column (LS column) that was attached to a magnetic stand (all from Miltenyi Biotec). After three rounds of washing, the column was removed from the magnetic stand and labelled cells were eluted in culture media for replating.

### Metabolic flux analyses using Seahorse XFe96 Analyzer

Mitochondrial oxygen consumption rates (OCR) and extracellular acidification rates (ECAR) were measured using a XFe96 Seahorse Biosciences Extracellular Flux Analyzer (Agilent Technologies). Purified motor neurons or neural progenitor cells were plated onto a Matrigel pre-coated Seahorse 96-well plate at 125,000 neurons per well 24 hours prior to the assay. Culture media were changed with 175ul of fresh Seahorse DMEM basal medium 45 minutes prior to the assay. Seahorse analyzer injection ports were filled with 1μM oligomycin, 1μM FCCP or 0.5μM each of rotenone and antimycin A for OCR. For ECAR, 10mM glucose, 1μM oligomycin and 50mM 2-DG were used, following the manufacturer’s instructions. Levels of OCR and ECAR were recorded, normalized and quantified based on manufacturer’s instructions.

### Small molecule treatments in MN cultures

NAM and C12 were reconstituted in water and DMSO respectively and diluted in media at the desired concentrations of 0.5 mM^23^ and 5 μM respectively. Motor neurons at day 27 were plated at 75,000 cells per well of a 96-well plate. Treatment with the respective small molecule began at day 28, for a total of 8 days. Biological triplicates were performed with a minimum of 5 technical replicates each.

### RNA interference in MN cultures

Motor neuron cultures at day 27 were dissociated with Accutase and seeded at 2 million cells per well in a 6-well plate. At day 28, non-targeting siRNA or siRNAs (Santa Cruz Technologies) were individually complexed with Lipofectamine RNAiMAX following manufacturer’s instructions and added to the motor neuron cultures. For each well, 10 pmol of siRNAs and 8μl of Lipofectamine RNAiMAX were used. Cells were either harvested for RNA and protein analyses, fixed for immunostaining or metabolic flux analysis were performed 3 days after siRNA transfection.

### RNA extraction and RT-qPCR

Cells were harvested in Trizol reagent for RNA extraction following manufacturer’s instructions. Purified RNA was converted to cDNA using the High-Capacity cDNA Reverse Transcription kit, and quantitative PCR (qPCR) was performed on the QuantStudio 5 Real-Time PCR System using PowerUp™ SYBR™ Green Master Mix (all from Applied Biosystems). Gene expressions were normalized to HPRT and ACTB expression unless otherwise stated. Primers used are listed in Supplementary Table S3.

### SDS-PAGE and Western blot

Protein lysates were resolved in 12% SDS-PAGE gels or 4-20% precast gels in Tris-Glycine-SDS buffer. Proteins were then transferred to a nitrocellulose membrane and blocked with 5% milk in TBST buffer. Primary antibodies were diluted in 5% milk and incubated with the membranes overnight at 4 °C. Primary antibodies used are listed in Supplementary Table S4. Membranes were washed thrice in TBST buffer. The corresponding horseradish peroxidase secondary antibodies (Life Technologies) were then diluted 1:5000 in 5% milk and incubated at room temperature for 90 minutes. Blots were washed thrice before exposing to ECL for imaging.

### Mitochondria isolation

Mitochondria from motor neuron cultures were isolated using the MACS technology (Miltenyi Biotec). Motor neurons culture were first Accutase and wash twice with PBS before resuspending in ice cold Lysis buffer. The cells were then homogenized with a dounce homogenizer (Pestle B) 15 strokes. The homogenate was diluted with 1X separation buffer based on manufacturer’s instructions. Anti-TOM22 were added to magnetically label the mitochondria before incubating at 4°C for 1 hour. The cells were then washed twice and filtered prior to loading into the separation column (LS column) that was attached to a magnetic stand (all from Miltenyi Biotec). After three rounds of washing, the column was removed from the magnetic stand and labelled mitochondria were eluted in storage buffer for downstream applications.

### Measurements of Complex I activity

Cell lysates were prepared using Complex I Enzyme Activity Microplate Assay Kit (Abcam), following the manufacturer’s instructions. Activity of Complex I were recorded, normalized and quantified based on manufacturer’s instructions.

### Immunostaining of cultured cells

Cells were fixed in 4% paraformaldehyde for 15 minutes, permeabilized in 0.1% Triton X-100 for 15 minutes and blocked in buffer containing 5% FBS and 1% BSA for an hour at room temperature. Primary antibodies (Table S3) were diluted in blocking buffer and incubated overnight at 4 °C. Cells were washed thrice in PBS. The respective secondary antibodies were diluted 1:1500 in blocking buffer and incubated at room temperature, in the dark, for 90 minutes. DAPI was used at 0.1 μg/ml to visualize cellular nuclei.

### Immunohistochemistry using ALS patient tissues

Lumbar spinal cord tissue sections were cut from blocks of paraffin embedded ALS tissue (n=4) and control tissue (n=4), obtained from UCSD CNS biorepository. Six μm-thick tissue sections were de-paraffinized with histology grade CitriSolv (twice for 15 minutes each), followed by a graded alcohol series (100, 90, 70, and 50% ethanol (vol/vol) for 3 min each), then washed in water (twice for 3 min). Endogenous peroxidase activity was then quenched in 0.6% hydrogen peroxide in methanol (vol/vol) for 15 min. After a 20 min permeabilization step in 1X PBS, 0.2% TritonX100, antigen retrieval was performed in a pressure cooker at 120 °C for 20 min in high pH solution (1% Tris-based). Sections were blocked with 2% Fetal Bovine Serum (vol/vol) and incubated with MnSOD (K68ac) antibody (1:100) overnight at 4 °C.

The following day after equilibration to room temperature, sections were washed three times in 1X PBS before 60 min room temperature incubation with 150◻l secondary antibody. Signals were detected via chromogenic reaction using NovaRed for 1-3 minutes per section until desired staining was achieved. Counterstaining was performed with hematoxylin for 10 seconds. Sections were dehydrated before adding cover slips.

### Image acquisition and image analysis

Images were acquired using the high content microscope Phenix (Perkin Elmer) using the 20x air objective. Image analyses including cell counts and intensity measurements were performed using Columbus (Perkin Elmer).

For primary neurites analysis, neuronal projections from the soma size was determined based on SMI-32 staining. For soma size analysis, cellular nuclei were identified by DAPI, and the cytoplasmic area surrounding the nucleus was determined based on SMI-32 staining. Soma size area was measured by image analysis software (ImageJ, NIH) based on the cytoplasmic area surrounding the nucleus, excluding neuronal projections.

For patient tissues, all slides were scanned with Hamamatsu Nanozoomer 2.0HT Slide Scanner at the UCSD Microscopy Core. Using NDP.view 2 viewing software, scanned slides were evaluated at 1X and 20X magnifications. All neurons were evaluated in both anterior horn sections from a total of four, non-sequential tissue sections per patient, in order to ensure no overlap in neurons. K68Ac expression patterns and intensity were determined for all neurons using Fiji. Color deconvolution was performed using “H DAB” as the defined vector. Neurons were measured and quantification performed using “Colour_2” representing the VectorRed signal without the background from the counterstain (Colour_1 is hematoxylin). The region of interest was determined for each neuron and “Mean gray value” was used to quantify intensity. In order to convert intensity to Optical Density (OD) the formula used was: OD = log (Max intensity/Mean intensity) for 8-bit images. The resulting OD quantified the average darkness of the image due to DAB signal (thus representing MnSOD-K68ac stain).

### Statistical Analyses

At least three biological replicates were performed for each experiment. Statistical analysis comparing two groups were performed by means of a two-tailed unpaired Student’s t test. P values lower than 0.05 were considered significant. All results are presented as mean ± standard deviation unless otherwise specified.

### Human Ethics Statement

All tissues were collected under HIPAA (Health Insurance Portability and Accountability Act of 1996) compliant IRB (Institutional Review Board) supervised consenting process. Patients are provided the option to donate their CNS tissue postmortem to the ALS biorepository and provide consent during life. All consent forms and other legal documentation is handled by clinical research support staff and kept de-identified from the basic research team. Each patient has separate clinical, CNS and fibroblast identifiers assigned to them in order to maintain patient autonomy and anonymity.

## ACKNOWLEDGEMENTS

This work is supported by Institute of Molecular and Cell Biology (A*STAR Research Entities) and the following grants to SY Ng: NRF-NRFF2018-03 (National Research Foundation Singapore), NMRC/OFYIRG/0011/2016 (National Medical Research Council, Singapore). This work is also partially supported by grants from National Natural Science Foundation of China to Y Fan: (grant numbers 81370766 and 81570101). We are grateful to Kevin Eggan (Harvard University) for the ALS iPSC lines 18a, 29d, 47a and 19f. We also thank Cedars-Sinai Medical Center’s David and Janet Polak Foundation Stem Cell Core Laboratory for providing the sporadic ALS iPSCs (CS14isALS-Tn16, CS51isALS-Tn3 and CS89isALS-Tn16). JHH, MMS and BXH are supported by the National University of Singapore Research Scholarships (PhD). We also thank Winanto, Yong Hui Koh, Li Yi Tan, Yu Ying Chew and Wai Kein Wong for their technical help in this project.

## AUTHOR CONTRIBUTIONS

SYN, BSS and JHH conceptualized and designed the study. JHH, MMS, SYN performed the experiments and analyzed the data. AT, ZJK, VJWL, BXH and YF performed some experiments. JR, YCL, BSS and SYN supervised the study. SYN and JHH wrote the manuscript. All authors have read and approved the manuscript.

## DECLARATION OF INTERESTS

The authors declare no competing interests.

