## Supplemental Figures for "ALS Motor Neurons Exhibit Hallmark Metabolic Defects That Are Rescued by Nicotinamide and SIRT3 Activation"

Correspondence to:

### Supplementary Tables

**Supplementary Table S1**: List of cell lines used

| **Cell Lines** | **Source** | **Catalog no.** |
|---|---|---|
| BJ-iPS iPSC | Ng et al. (2015) | N/A |
| 18a iPSC | Eggan Lab, Boulting et al. (2011) | N/A |
| GM23720 iPSC | Coriell Institute | GM23720 |
| 29d iPSC ($SOD1^{L144F}$) | Eggan Lab, Boulting et al. (2011) | N/A |
| 47a iPSC ($TDP43^{G298S}$) | Eggan Lab, Rodriguez-Muela et al. (2017) | N/A |
| 19f iPSC (C9ORF72) | Eggan Lab, Kiskinis et al. (2014) | N/A |
| CS14isALS-Tn16 (sALS1) iPSC | Cedars-Sinai Medical Center's | CS14isALS-Tn16 |
| CS51isALS-Tn3 (sALS2) iPSC | Cedars-Sinai Medical Center's | CS51isALS-Tn3 |
| CS89isALS-Tn16 (sALS3) iPSC | Cedars-Sinai Medical Center's | CS89isALS-Tn16 |
| BJ-$SOD1^{L144F}$ iPSC | This paper | N/A |
| BJ-$TDP43^{G298S}$ iPSC | This paper | N/A |
| BJ-SIRT3+/- #6 iPSC | This paper | N/A |
| BJ-SIRT3+/- #17 iPSC | This paper | N/A |

**Supplementary Table S2**: List of oligonucleotides used for CRISPR/Cas9 studies

| **Oligonucleotides (CRISPR)** | **Source** |
|---|---|
| SOD1$^{L144F}$ sgRNA (CACCGAGGAAACGCTGGAAGTCGTT) | Integrated DNA Technologies |
| SOD1$^{L144F}$ ssODN (ACATCCAAGGGAATGTTTATTGGGCGATCCCAATTACACCA CAAGCGAAACGACTTCCAGCGTTTCCTGTCTTTGTACTTTCTT CATTTCCACCTTTGCC) | Integrated DNA Technologies |
| SOD1$^{L144F}$ surveyor F: (TAAGGGTAGCGTGTGGTGGT) SOD1$^{L144F}$ surveyor R: (TGCTTAGACAAATAGGCTGTCC) | Integrated DNA Technologies |
| TDP43$^{G298S}$ sgRNA (CACCGTTTGGTAATAGCAGAGGGGG) | Integrated DNA Technologies |
| TDP43$^{G298S}$ ssODN (TTTGGGAATCAGGGTGGATTTGGTAATAGCAGAGGGGGTGG AGCTGGTTTGGGAAACAATCAAGGTAGTAATATGGGTGGTG GGATGAACT) | Integrated DNA Technologies |
| TDP43$^{G298S}$ surveyor F (CCACTACGCCCAGCTAATGT) | Integrated DNA Technologies |
| TDP43$^{G298S}$ surveyor F (TCTGGCTGGGGAATGTAGAC) | Integrated DNA Technologies |
| SIRT3$^{+/-}$ sgRNA (CTTCCGGCGCCGAGCGGCGCGG) | Integrated DNA Technologies |
| SIRT3$^{+/-}$ surveyor F: (GGCGCTCACTTCTTCGTGTA) SIRT3$^{+/-}$ surveyor R: (AGACGTAGAGGCGAGTAGAGGA) | Integrated DNA Technologies |

**Supplementary Table S3**: List of human primers used in qPCR studies

| **Oligonucleotides (qPCR)** | **Source** |
|---|---|
| CHOP qPCR F:<br>(AAGGCACTGAGCGTATCATGT)<br>CHOP qPCR R:<br>(TGAAGATACACTTCCTTCTTGAACA) | Ng et al. (2015) |
| sXBP1 qPCR F:<br>(TGCTGAGTCCGCAGCAGGTG)<br>sXBP1 qPCR R:<br>(GCTGGCAGGCTCTGGGGAAG) | Ng et al. (2015) |
| ACTB qPCR F:<br>(CCAACCGCGAGAAGATGA)<br>ACTB qPCR R:<br>(CCAGAGGCGTACAGGGATAG) | Ng et al. (2015) |
| HPRT qPCR F:<br>(TATGGCGACCCGCAGCCCT)<br>HPRT qPCR R:<br>(CATCTCGAGCAAGACGTTCAG) | Integrated DNA Technologies |

**Supplementary Table S4**: List of antibodies used in western blot and immunostaining studies

| **Antibodies** | **Company** | **Catalog no.** |
|---|---|---|
| Rabbit anti-SirT3 (D22A3) | Cell Signaling | 5490 |
| Mouse anti-alpha tubulin (B-7) | Santa Cruz | sc-5286 |
| Mouse Anti-TOMM20 | Abcam | ab56783 |
| Rabbit anti-Islet 1 [EP4182] | Abcam | ab109517 |
| Mouse anti-SMI-32 | BioLegend | 801701 |
| Rabbit anti-Acetylated-Lysine | Cell Signaling | 9814 |
| Rabbit anti-SOD2/MnSOD | Abcam | ab13533 |
| Rabbit anti- SOD2/MnSOD (acetyl K68) | Abcam | ab137037 |
| Rabbit anti-SOX1 | Abcam | ab87775 |
| Mouse anti-NESTIN [10C2] | Abcam | ab22035 |
| Mouse anti-Troponin T | Thermo Fisher Scientific | MS-295-P |
| CD171 (L1CAM)-APC | Miltenyi Biotec | 130-100-684 |
| Anti-PSA-NCAM-APC | Miltenyi Biotec | 130-120-437 |
| AlexaFluor Donkey anti-Mouse 488 | Thermo Fisher Scientific | A21202 |
| AlexaFluor Donkey anti-Rabbit 647 | Thermo Fisher Scientific | A31573 |
| AlexaFluor Donkey anti-Rabbit 488 | Thermo Fisher Scientific | A21206 |
| AlexaFluor Donkey anti-Mouse 568 | Thermo Fisher Scientific | A10037 |
| Goat anti-rabbit IgG, HRP conjugated | Thermo Fisher Scientific | 31466 |
| Goat anti-mouse IgG, HRP conjugated | Thermo Fisher Scientific | 31431 |
